# RNA polymerase II and PARP1 shape enhancer-promoter contacts

**DOI:** 10.1101/2022.07.07.499190

**Authors:** Gilad Barshad, James J. Lewis, Alexandra G. Chivu, Abderhman Abuhashem, Nils Krietenstein, Edward J. Rice, Oliver J. Rando, Anna-Katerina Hadjantonakis, Charles G. Danko

## Abstract

How enhancers control target gene expression over long genomic distances remains an important unsolved problem. Here we studied enhancer-promoter contact architecture and communication by integrating data from nucleosome-resolution genomic contact maps, nascent transcription, and perturbations to transcription-associated proteins and thousands of candidate enhancers. Contact frequency between functionally validated enhancer-promoter pairs was most enriched near the +1 and +2 nucleosomes at enhancers and target promoters, indicating that functional enhancer-promoter pairs spend time in close physical proximity. Blocking RNA polymerase II (Pol II) caused major disruptions to enhancer-promoter contacts. Paused Pol II occupancy and the enzymatic activity of poly (ADP-ribose) polymerase 1 (PARP1) stabilized enhancer-promoter contacts. Based on our findings, we propose an updated model that couples transcriptional dynamics and enhancer-promoter communication.

## Main Text

Much of metazoan cellular diversity is encoded by *cis*-regulatory elements known as enhancers, which regulate the rate of mRNA production from distal promoters (*1*). Since the landmark discovery of the SV40 enhancer more than 40 years ago (*2*–*4*) a key goal has been to understand the molecular basis by which enhancers and promoters communicate across long stretches of DNA sequence. The predominant model proposes that enhancers and promoters ‘loop’ into close physical proximity in the 3D space within the nucleus. These loop interactions are frequently represented as a physical bridge by which the enhancer and promoter are connected via highly stereotyped protein-protein interactions involving transcription factors, Pol II, mediator, cohesin and other proteins (*5, 6*). Indeed, chromosome conformation capture (3C) based methods such as *in-situ* Hi-C (*7*) and Micro-C (*8*), which measure the frequency of ligation between DNA sequences that are close together in 3D space (*9*), can be used to predict the functional impact of enhancers on a target genes (*10*–*12*). Moreover, changes in enhancer-promoter loops (*13, 14*), as well as preestablished loops (*10, 15*) are associated with activation of target promoters.

Despite some support, however, several recent observations are not compatible with the looping model. Measurements of enhancer-promoter distances in several fly and mouse developmental loci, using microscopy in both living and fixed cells, have revealed that enhancers and promoters are hundreds of nanometers apart at the time of gene activation (*16*–*18*). In one well-characterized locus, the physical distance between the *Shh* promoter and several developmental enhancers increased following gene activation (*16*). Finally, depletion of proteins proposed to constitute a physical bridge, such as mediator and cohesin, have minimal impact on either Hi-C maps or transcription (*19, 20*). These studies have demonstrated that we still lack complete answers to long-standing questions about enhancer-promoter communication: Do active enhancers “linger” close to their target promoters in order to activate transcription? And which molecules play a role in facilitating enhancer-promoter communication?

Here we leverage the new high-resolution 3C method, Micro-C (*8, 21*–*23*), nascent RNA sequencing (*24*–*26*), and perturbations to Pol II, PARP1, and thousands of candidate enhancers (*27, 28*), to study the interplay between transcription and enhancer-promoter contact dynamics. We used CRISPR interference (CRISPRi) experiments testing nearly six thousand candidate enhancers (*27, 28*) to demonstrate that functional interactions between enhancers and their target promoter were associated with increased Micro-C contact frequency compared with those lacking an impact on gene expression, suggesting that functional enhancer-promoter pairs spend more time in very close physical proximity. Manipulation of transcription-related proteins revealed a crucial role for Pol II and its transcriptional dynamics affecting enhancer-promoter contact frequency. We also found that the enzymatic activity of poly (ADP-ribose) polymerase 1 (PARP1), which creates a web of post-translational modifications and transient interactions via PAR chains that connect transcription and chromatin proteins, had a stabilizing effect on enhancer-promoter contacts. These observations lead us to an updated model that integrates the effects of transcription and chromatin in enhancer-promoter communication.

## Results

### Enhancer function correlates with enhancer-promoter contacts

We asked whether functional enhancer-promoter pairs, in which enhancers are actively stimulating gene expression, spend more time in close physical proximity compared to nonfunctional pairs. To detect interactions between candidate enhancers and promoters, we generated a ∼1.7 billion contact Micro-C dataset in K562 cells (**Fig. 1A**). We defined enhancer function as the effect an enhancer has on target gene expression or activity using CRISPRi to knock down candidate enhancers (*11, 27*–*29*).

**Figure 1.**
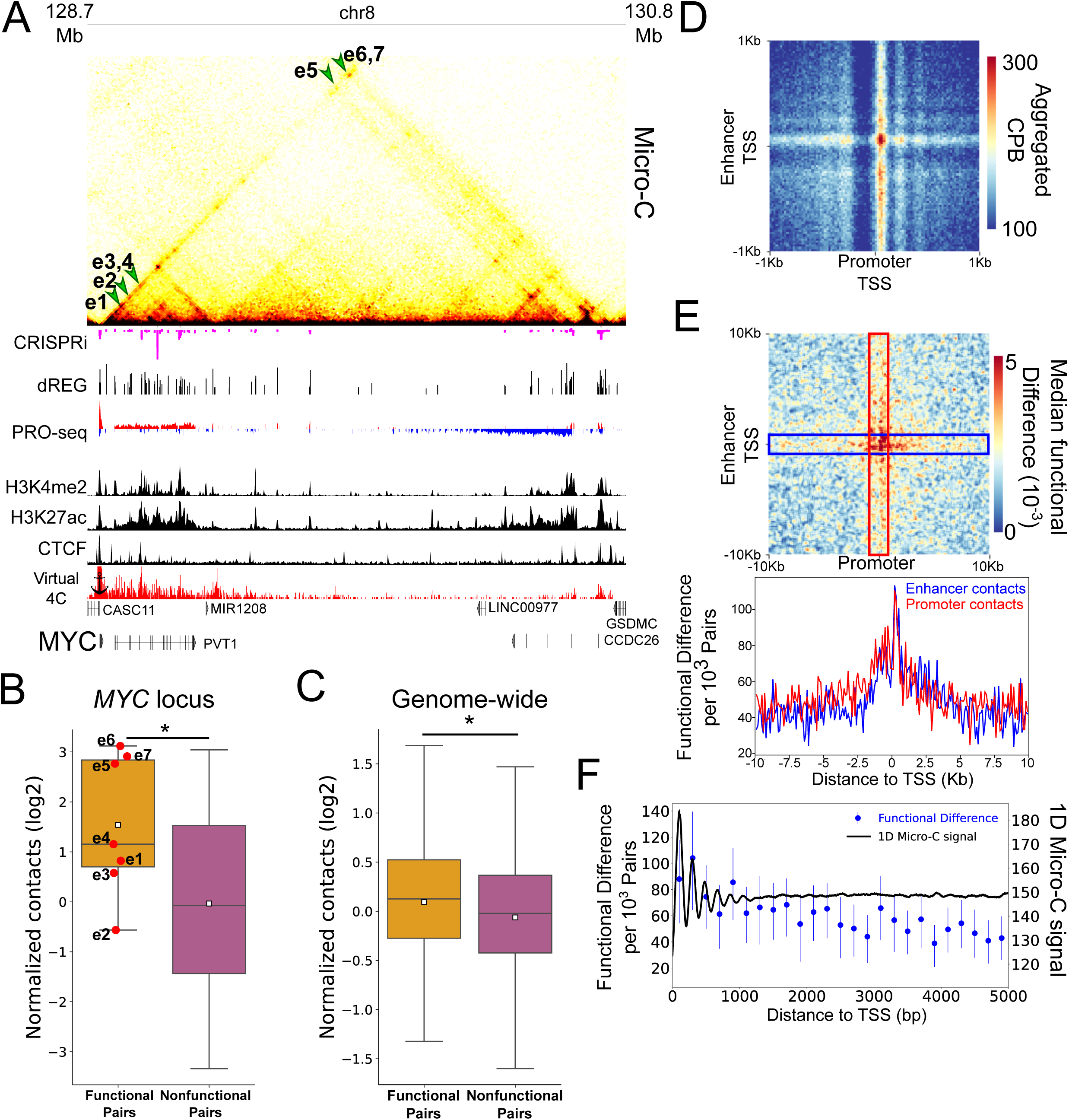
Micro-C contacts are enriched in functional enhancer-promoter pairs. (A) Genome browser tracks showing Micro-C contact maps, CRISPRi-associated changes in cell viability, dREG-defined TIRs and input PRO-seq signal, H3K4me2, H3K27ac and CTCF ChIP-seq signal and a 2D representation (virtual 4C) of Micro-C contacts with the *MYC* promoter, along the *MYC* locus. Green arrows point to the regions in the contacts map representing interaction between the seven CRISPRi-validated *MYC* enhancers and the *MYC* promoter (B) Box-plot comparing the observed contact frequency relative to expected by a local distance-decay function of the seven validated functional enhancers in the *MYC* locus (functional pairs) compared to the rest of the dREG-detected TIRs in the TAD (nonfunctional pairs) with the *MYC* promoter. (C) Box-plot comparing contact levels relative to expected by a local distance-decay function of functional versus the nonfunctional enhancer-promoter pairs in the genome, matched for genomic distance, accessibility and target gene expression distributions. (* Mann-Whitney p-value < 0.05). (D) An APA for enhancer-promoter contacts at 2kb around the TSSs. Pixel size is 20bp square and color scale represents contacts per billion contacts (CPB). (E) Top panel - APA for the differences in contacts between functional and nonfunctional pairs based on CRISPRi. The APA represents smoothed contact differences at a region of 20kb around the TSS with original pixel size of 100bp square. Red and blue rectangles represent the regions where contacts are plotted at the bottom panel. Bottom panel - metaplot of the differences in contacts between functional and nonfunctional pairs in a 2kb window around the TSS (the TSS region). As indicated in the red and blue rectangles over the APA map, at the metaplot the red line represents the differences in contacts between functional and nonfunctional pairs associated with the promoter TSS region and 20kb around enhancer TSSs (promoter contacts). The blue line represents the differences in contacts between functional and nonfunctional pairs associated with the enhancer TSS region and 20kb around promoter TSSs (enhancer contacts). (F) Blue - dot and error plot showing the median difference between functional and nonfunctional pairs (dot) and the associated 95% confidence interval of the median based on 1000 bootstrap iterations (error-bars) in enhancer-promoter contacts associated with enhancers’ and promoters’ +1 and +2 nucleosomes (0-400bp downstream to the TSS). Contact differences in windows of 200 bp from the TSS are shown at the center coordinate for each window. Black - one dimensional contact signal downstream to enhancers and promoters. Signal is calculated for intra-chromosomal contacts with both sides having mapping quality (mapq) > 30. Total median signal was smoothed using a sliding window of 100bp.

We first analyzed seven CRISPRi-defined enhancers that regulate *MYC* expression in K562 cells (*28*), all of which fall within the same topologically associated domain (TAD) as the *MYC* protein-coding gene. Each of the seven active *MYC* enhancers was located near a transcription initiation region (TIR), representing the transcription start site of an enhancer RNA (eRNA), identified from dREG (*30, 31*) analysis of K562 PRO-seq data (*32*) (**Fig. 1A**). We calculated the number of contacts observed between each TIR and the *MYC* promoter, normalized for the local background and the linear distance between each enhancer-promoter pair (**Fig. S1A; see Methods**). Contacts between the seven CRISPRi-validated enhancers and the *MYC* promoter (functional enhancer-promoter pairs) were significantly higher than those observed for the other TIRs within the same TAD that had no detectable effect on *MYC* (nonfunctional pairs, ∼2.46 fold increase in median normalized contacts for functional pairs, *p* = 0.015; Mann-Whitney U-test) (**Fig. 1B**).

As the *MYC* locus in K562 cells is unusual in many respects, we next examined the relationship between enhancer function and enhancer-promoter contacts genome-wide. We used data from a CRISPRi screen that tested the function of 5,920 candidate enhancers in K562 cells (*27*). We defined 620 functional enhancers that are located between 5kb and 1Mb away from their target promoter and that elicited a significant decrease in target gene expression following CRISPRi knockdown (empirical p-value < 0.05 and expression fold change < 0.9). As a control, we defined 3,375 nonfunctional pairs in which the enhancer had no effect on a particular target gene (empirical p-value > 0.9 and expression fold change within 0.95-1.05). We found a significantly higher number of contacts in functional enhancer-promoter pairs (∼17% increase in median normalized contacts for functional pairs, *p* < 1×10^−10^; Mann Whitney U test) (**Fig. S1B**). Next we curated a subset of 196 functional and 196 nonfunctional enhancer-promoter pairs, in which we controlled for the confounding effects of linear distance, target gene transcription levels, and chromatin accessibility (**Fig. S1C**). Here too, we found a significantly higher number of contacts in functional enhancer-promoter pairs (∼12% increase in median normalized contacts for functional pairs, *p* = 0.028; Mann Whitney U test) (**Fig. 1C**). We conclude that functionally active enhancer-promoter pairs, in which the enhancer is actively controlling expression from the target promoter, have a higher interaction frequency in Micro-C data than nonfunctional pairs.

We next investigated whether functional enhancer-promoter pairs spend more time close together in 3D space. Micro-C requires chromatin to be close enough to crosslink in a confirmation and at a distance that the free ends of DNA exiting each nucleosome can ligate to one another (*9*). We reasoned that if enhancer-promoter pairs are predominantly ∼100-200 nm apart during communication, as observed in live and fixed cell imaging studies (*16*– *18*), increased Micro-C signal between functional pairs will be most prominent further from the transcription start site (TSS) where chromatin can reach out through the intervening nuclear space, crosslink and ligate. By contrast, increased signal near the TSS at both the enhancer and promoter implies that enhancer-promoter pairs can be close enough that crosslinked chromatin can swing around and ligate directly to one another. We therefore examined the enrichment of functional contacts as a function of distance from the TSS. Active enhancers and promoters consist of a nucleosome-free core region flanked by bidirectional transcription initiation and well-positioned +1 and +2 nucleosomes downstream of the most-used TSS (*33*–*35*) that are readily observed in Micro-C data after aligning on coPRO TSSs (*32, 36*) (**Fig. S2**). Aggregated peak analysis (APA) between all candidate enhancer and promoter pairs (5kb-100kb) showed that contacts between +1 (promoter)/ +1 (enhancer) nucleosomes were most prominent (**Fig. 1D**). Next, we examined the difference in contact frequency between the CRISPRi functional and nonfunctional enhancer promoter pairs. The difference was highest near the TSS, especially involving the +1 and +2 nucleosomes, and decayed as a function of distance to ∼5kb from the TSS (Pearson’s R = -0.81, *p* = 9.6×10^−7^) (**Fig. 1E,F**). We conclude that contact frequency between functional enhancer-promoter pairs was most enriched within the first ∼1 kb from the TSS. This result indicates that enhancer activation of a target promoter was associated with a relatively high probability of the enhancer-promoter pair being in close proximity in 3D space.

### Enhancer-promoter contacts depend on active transcription

We asked which cellular factors mediate the increased contacts observed between enhancers and their target promoters. One model of interaction involves the aggregation of transcription proteins into clusters that contain both enhancers and promoters and act to facilitate communication (*37*–*43*). Both the C-terminal domain (CTD) of the large subunit of RNA Pol II and nascent RNA are reported to form macromolecular clusters with other transcription-related proteins (*40, 44, 45*). These results imply that Pol II itself may play a role in mediating enhancer-promoter contacts. Although perturbing Pol II was reported to have modest effects on enhancer-promoter contacts (*22, 46*), previous studies have generally not accounted for global changes in the proportion of the entire Micro-C library (i.e., 1D signal) that map to enhancer- or promoter-regions, and thereby may have underestimated the effects of Pol II depletion.

We set out to test the hypothesis that Pol II is required for enhancer and promoter regions to come into close proximity. To accommodate global changes in the distribution of contacts, we devised APAs that adjust for local background near enhancer- and promoter-anchors between different treatment conditions (**Fig. 2A; see Methods**). Using this strategy to re-analyze Micro-C data after blocking either Pol II initiation (triptolide - TRP) or release from pause (flavopiridol - FLV) (*22*) showed that the largest effect of Pol II transcriptional inhibition occurred near the TSS (**Fig. 2B**), consistent with the hypothesis that actively elongating Pol II plays a pivotal role in holding enhancers and promoters in physical proximity. Moreover, by blocking release from pause, FLV not only prevents actively elongating Pol II from entering the gene body, but also leaves paused Pol II near the TSS at most promoters (*47*). We hypothesized that the presence of paused Pol II may retain some of the interactions that are depleted in TRP, in which all Pol II is depleted from chromatin. Indeed, inhibition of Pol II recruitment to promoters and enhancers by TRP had a stronger effect on enhancer-promoter contacts compared with the effect of inhibiting pause release by FLV (**Fig. 2B, lower panel**; *p* < 10^−100^; Wilcoxon signed-rank test), suggesting that Pol II occupancy at the pause site may have a stabilizing effect on these contacts.

**Figure 2.**
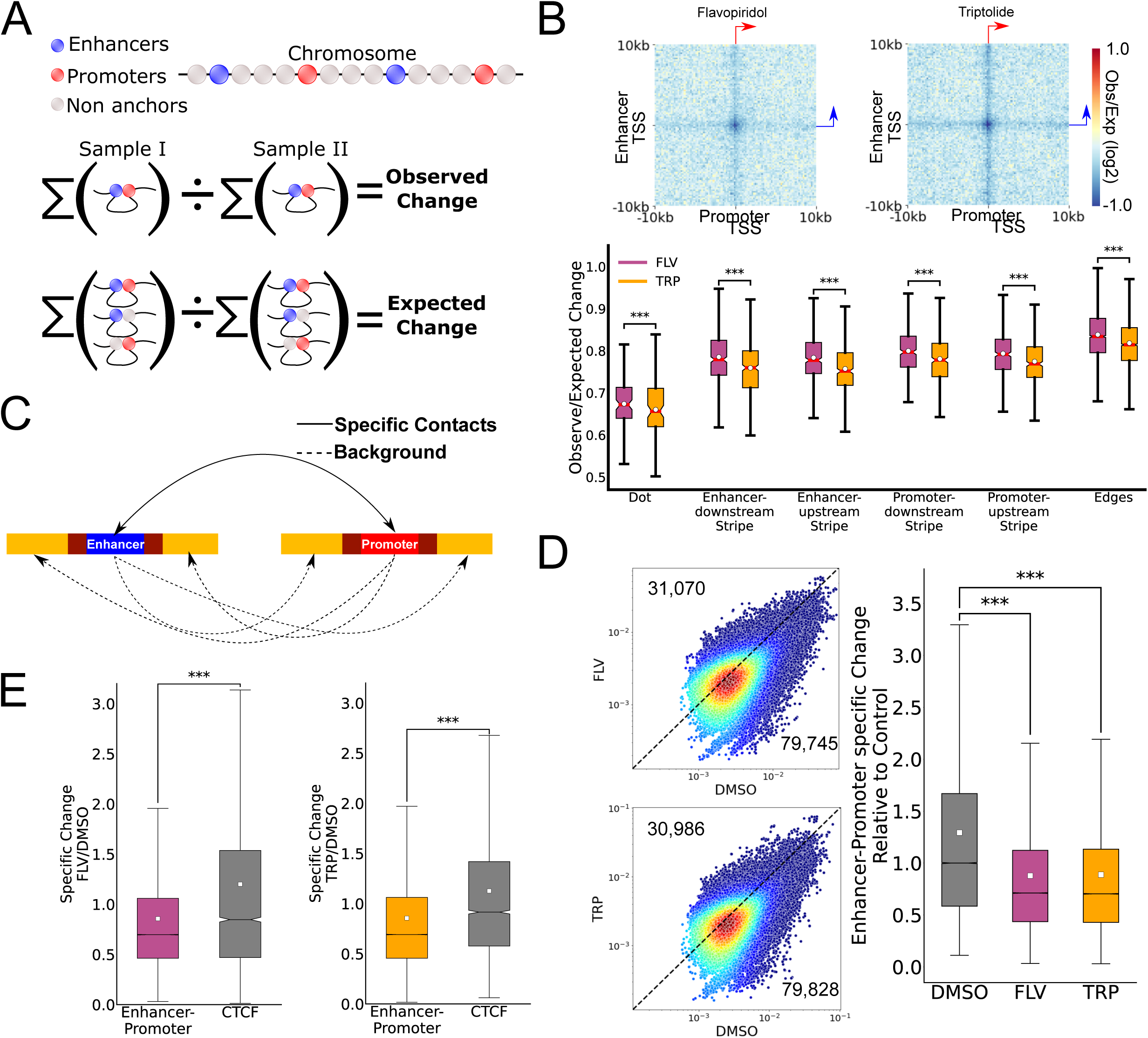
Enhancer-promoter contacts depend on active transcription. (A) Schematic representation of the strategy used to compute APAs comparing between different conditions. (B) APA heatmap representation of the change in contacts compared to DMSO control for mESCs treated with FLV or TRP (top) and box-plots showing the observed over the expected change at the dot, the different stripes and the edges of the APA change matrices in each of the TRP and FLV treatments (*** Wilcoxon signed-rank test p-value < 1×10^−100^). (C) Schematic representation of ratio calculation between enhancer-promoter contacts (solid arrows) and background contacts (dashed arrows). (D) Scatterplot comparing the enhancer-promoter contacts over background ratio between FLV-treated (top) or TRP-treated (bottom) and DMSO-treated (control) mESCs. Box-plots show the ratio distribution relative to the median control (DMSO) ratio. (E) Boxplots showing the ratio distribution relative to the median control (DMSO) ratio for enhancer-promoter pairs and transcriptionally inactive bound CTCF motifs (*** Mann-Whitney p-value < 1×10^−100^).

To complement the normalization performed in APAs, we also devised an alternative local normalization scheme that is specific to each candidate enhancer-promoter pair (**Fig 2C; see Methods**). Consistent with the APA analysis, we found that following transcription inhibition enhancer-promoter contacts dropped by ∼30% (**Fig. 2D**). Additionally, more than 70% of enhancer-promoter pairs showed lower contact frequency following acute transcriptional inhibition (**Fig. 2D**). To explore the specificity of the effect of Pol II inhibition on enhancer-promoter contacts, we analyzed its effect on CTCF-associated contacts. CTCF binding sites often exhibit strong focal contacts in Hi-C and Micro-C maps (*21*) but did not show the same decreased contact frequency observed between enhancer-promoter pairs (**Fig. 2E**). We conclude that focal contacts between enhancers and promoters, but not CTCF-CTCF contacts, are substantially depleted following acute transcriptional inhibition, indicating a role for Pol II in facilitating contacts.

### Rates of Pol II initiation and pause correlate with enhancer-promoter contacts

We next asked how different steps in the transcription cycle correlate with contacts. At steady-state, the rate of transcription initiation is proportional to gene body transcription levels, whereas the rate of release of paused Pol II into productive elongation is proportional to the pausing index (*48*). We divided human gene promoters into 4 quartiles based on their gene body transcription levels (initiation), the gene body-normalized PRO-seq signal at the first 250bp downstream of the TSS (pausing index) or the PRO-seq signal at the first 250bp downstream of the TSS alone (pausing signal) in K562 cells (**Fig 3A,B**). The largest change in overall promoter contacts with the surrounding enhancers was associated with gene body transcription levels, in-line with previous findings (*11, 14, 49*). Increased enhancer-promoter contacts were also associated with pausing signal and pausing index. However, whereas the increase in contacts associated with gene body transcription spread across the regions surrounding enhancers and promoters, the pause-associated correlation was more specific to focal (TSS-TSS) enhancer-promoter contacts near the location at which paused Pol II resides (**Fig 3B, S3**).

**Figure 3.**
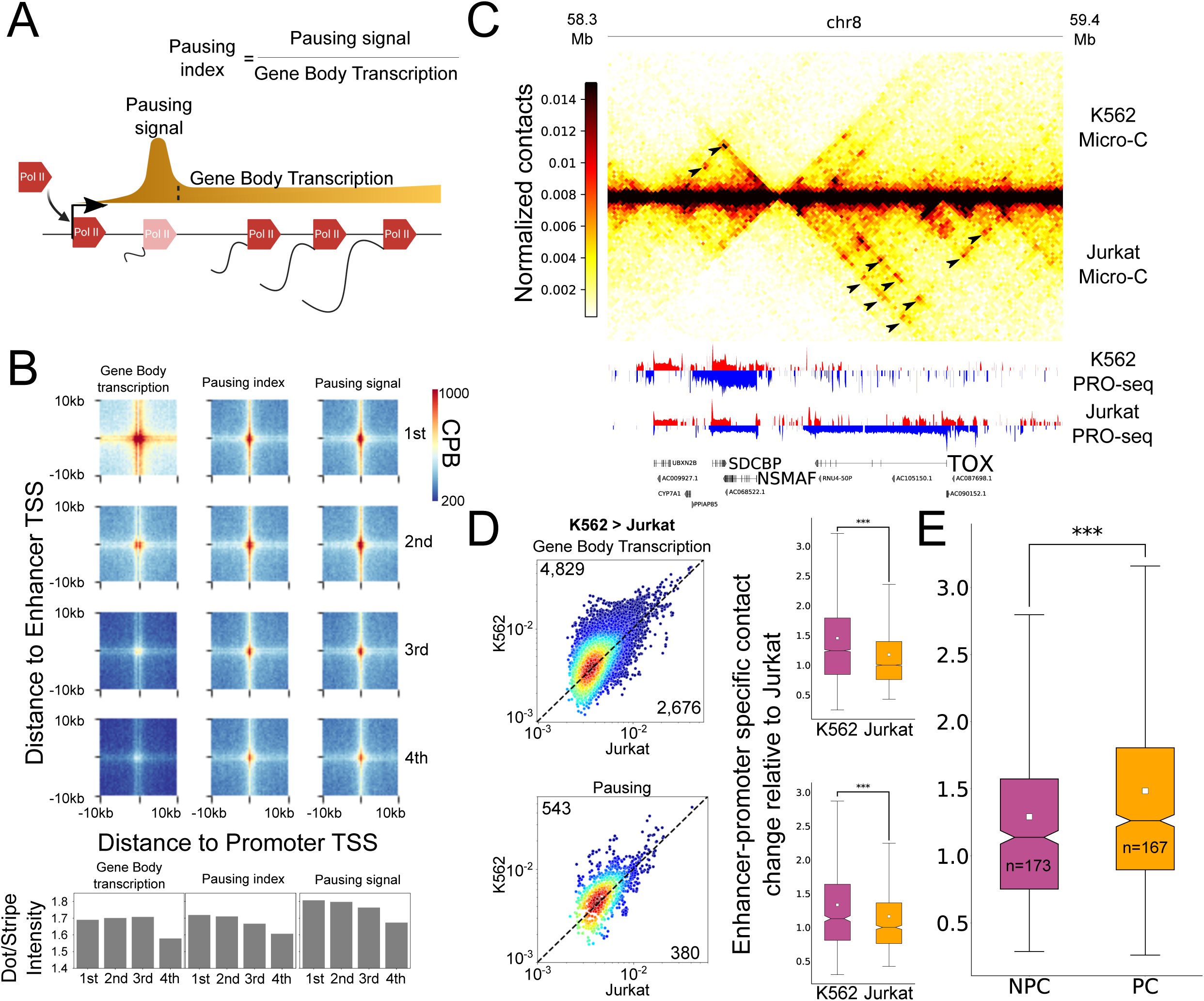
Changes in Pol II pausing and gene body density correlate with enhancer-promoter contacts. (A) Schematic representation of gene transcription from initiation, pausing and productive elongation at the gene body. The definitions for the PRO-seq signal at the pause peak (250bp downstream to the TSS - pausing signal) and gene body, as well as the calculated pausing index for this analysis, are illustrated. (B) APA maps show enhancer-promoter contacts associated with promoters of the four gene body transcription (left), pausing index (middle) and pause peak signal (right) quartiles. The color scale represents contacts per billion contacts (CPB) and bar-plots demonstrate the ratios between the dot- and stripe-associated contacts in the APAs. (C) Genome-browser shot of a 1.1Mb region containing the *TOX* and *NSMAF* genes. The Micro-C contact map pixel size is 10kb. Arrows indicate differential contacts associated with differential transcriptional activity in K562 and Jurkat T-cells. (D) Scatter plots and boxplots examine how changes in enhancer-promoter contacts are associated with promoters of genes with significantly higher gene body transcription (top) and pausing signal (bottom) at K562 compared to Jurkat (*** Wilcoxon signed-rank test p-value < 1×10^−100^). (E) Boxplot shows the relative increase of enhancer-promoter contacts associated with promoters of genes with upregulated gene body transcription in K562 with a corresponding significant increase in pausing signal (pause change - PC) and without a change in pausing signal (no pause change - NPC). (*** Mann-Whitney p-value < 1×10^−100^)

To further isolate the effect of Pol II pausing from productive elongation, we compared changes in transcription and contacts between different cell types. We generated new Micro-C data from Jurkat T-cells (∼1.18 billion contacts) and compared it to our K562 Micro-C data. Both Jurkat and K562 are cultured cell lines of human origin, but they model different cell types in the hematopoietic lineage; While K562 show similar properties to cells of the common myeloid progenitor lineage, Jurkats model human T-cells. Overall, transcriptional differences between the cell lines were associated with differences in enhancer-promoter contacts (**Fig 3C**). Differential transcription of gene bodies, and differences in the abundance of paused Pol II near promoters, were both positively correlated with enhancer-promoter contacts (**Fig 3D, Fig. S4A**). To isolate the effects of paused Pol II on contact frequency, we compared gene promoters associated with a similar and significant change in gene body transcription, which differ because they were either associated with a change or no associated change in paused Pol II levels (**Fig. S4B**; PC = Pause change; NPC = No pause change). We found that genes with a significant increase in productive elongation but no associated change in paused Pol II showed either no increase or a smaller increase in enhancer-promoter contacts, relative to genes associated with increased paused Pol II (**Fig. 3E, Fig. S4C**). Hence, we conclude that paused Pol II has a significant effect on enhancer-promoter contacts that is independent of initiation or productive elongation rates.

### NELF degradation depletes enhancer-promoter contacts

To further explore the role of Pol II pausing, we next asked whether depleting paused Pol II removed enhancer-promoter contacts. Although previously published triptolide and flavopiridol experiments alter Pol II pausing, they also appear to have a substantial inhibitory effect on transcription initiation (*50*). To focus on the effect of Pol II pausing, we used a mouse embryonic stem cell (mESC) system in which both copies of the negative elongation factor complex submit B (NELFB) were tagged with FKBP12^F36V^, allowing the rapid and reversible degradation of the NELF complex in the presence of a dTAG ligand (*51, 52*) (**Fig. 4A**). Following 30 minutes of NELFB depletion, Pol II density in TSSs decreased. However, by 60 minutes of NELFB depletion, Pol II signal near the TSS was partially regained (**Fig. 4B**). Notably, it was recently shown that this recovery of Pol II near the TSS represents transcriptionally inactive Pol II that cannot productively elongate in the absence of NELF (*53, 54*). This suggests that while Pol II pausing was depleted following NELFB depletion, transcription initiation rates were intact or may even increase (*50*).

**Figure 4.**
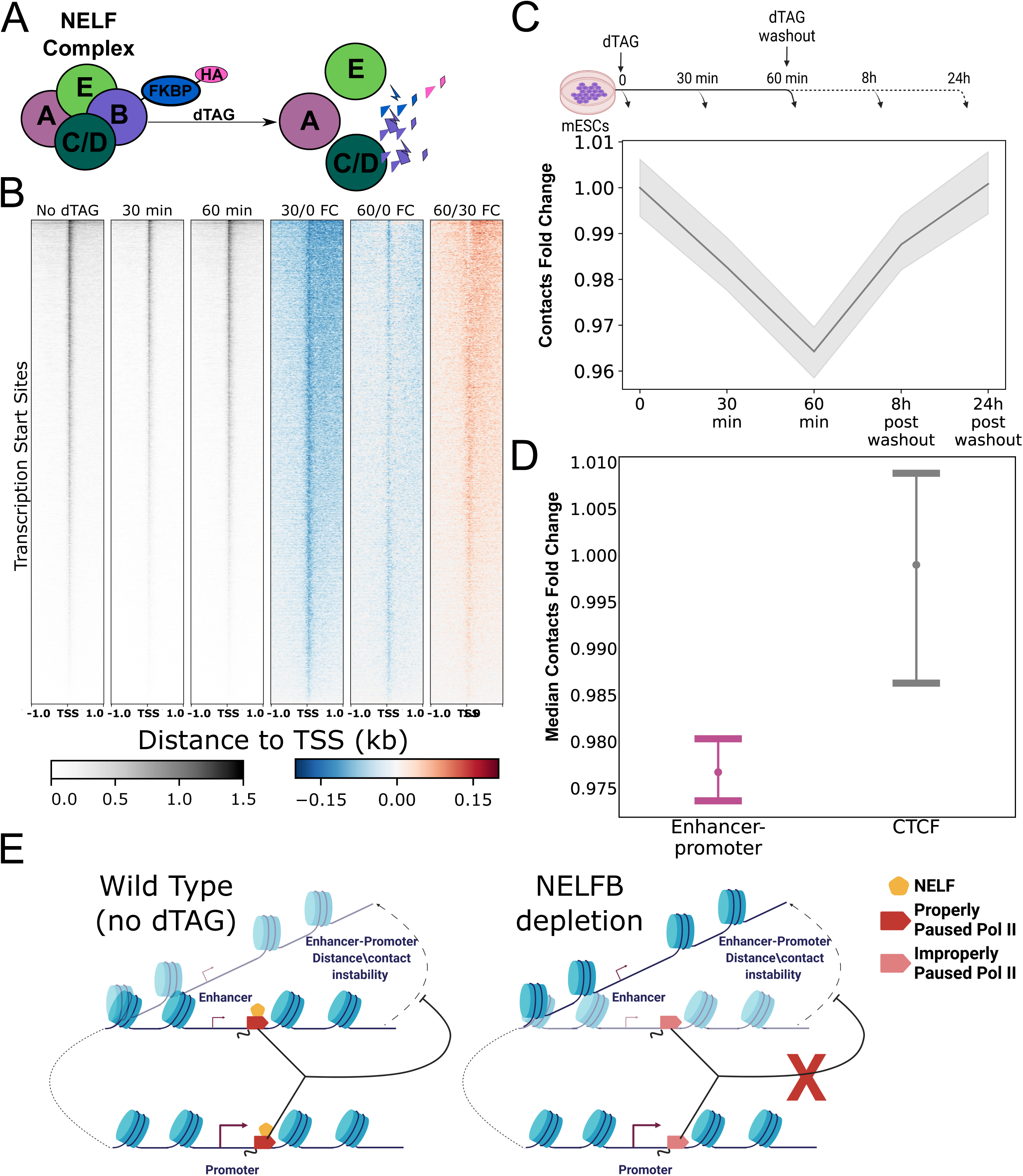
NELFB depletion and recovery correlates with changes in enhancer-promoter contacts. (A) Illustration of the NELFB-dTAG system and the corresponding effect on the NELF complex. Following NELFB depletion with dTAG, the NELF complex dissociates and is no longer found bound to chromatin (*93*) (B) Heatmaps showing PRO-seq signal in untreated (control) mESCs (left), the fold-change in PRO-seq signal following 30 minutes of dTAG treatment and after 60 minutes of dTAG treatment near TSSs as well as the fold change in PRO-seq signal at 30 minutes and 60 minutes of dTAG treatment compared to untreated control and at 60 minutes compared to 30 minutes of dTAG treatment. (C) A line plot showing the median enhancer-promoter contacts over background ratio change relative to the untreated (T=0) control. Gray shadow represents the 95% confidence interval for the median, based on 1000 bootstrap iterations. (D) Dot and error plot demonstrating the median contact change and the 95% confidence interval of the median based on 1000 bootstrap iterations, after 60 minutes of NELFB depletion, for enhancer-promoter (purple) and transcriptionally inactive CTCF motif contacts (gray). (E) Model shows the effect of properly paused (in the presence of NELF) and improperly paused Pol II on enhancer-promoter contacts.

To ask if such a drop in Pol II pausing results in a loss of enhancer-promoter contacts, we generated Micro-C libraries (∼300 million contacts each) following a time-course of NELFB depletion and dTAG washout. We found a small but highly reproducible drop in enhancer-promoter contacts beginning at 30 minutes and decreased further at 60 minutes of NELFB depletion (**Fig. 4C; Fig. S6**). This suggests that the accumulation of improperly paused Pol II (*53, 54*) cannot rescue the loss of contacts associated with paused Pol II loss. Washout of the dTAG ligand over 8 and 24 hours, corresponding to a ∼20-40% restoration of NELFB levels (*54*), increased enhancer-promoter contacts back to the levels observed in untreated cells (**Fig. 4C; Fig. S6**). The effect of dTAG was specific to enhancer-promoter contacts and was not observed at transcriptionally inactive CTCF binding sites (**Fig 4D**). Finally, the magnitude of decrease in contact frequency correlated with the magnitude of paused Pol II loss at 30 minutes (Pearson’s R = 0.24; *p* = 0.018; taking median of 1-percentiles, see Methods), such that candidate enhancer-promoter pairs which lost more paused Pol II also lost more contacts. A good example is the ZRS enhancer of the *Shh* gene, which had a large drop in paused Pol II signal as well as a large loss of contacts with the *Shh* promoter following 30 minutes of NELFB depletion (**Fig. S5**). Hence, we conclude that paused Pol II contributes to enhancer-promoter contact levels (**Fig 4E**).

### PARP1 enzymatic activity has two opposite effects on enhancer-promoter contact stability

The effect of paused Pol II highlighted another protein known to be involved in interactions between transcription-related proteins, PARP1, as a potential regulator of enhancer-promoter contacts. PARP1 is an enzyme which catalyzes the formation of poly (ADP-ribose) (PAR) chains, a post-translational modification added to transcription and chromatin-associated proteins (which are said to be “PARylated”) near active enhancer and promoter regions (*55*–*61*). PAR chains play a critical role in driving the formation of molecular clusters through interactions with intrinsically disordered regions (IDRs) of proteins (*62*). Finally, inhibiting the enzymatic activity of PARP1 prevented changes in 3D distance between the *Shh* promoter and developmental enhancers in mouse embryos (*16*).

We asked whether PARP1 enzymatic activity impacts enhancer-promoter contacts. We inhibited PARP1 enzymatic activity using the small molecule inhibitor Olaparib, which is relatively specific to PARP1 and leaves catalytically inactive PARP1 on chromatin (*63*). We generated Micro-C libraries following 90 minutes of Olaparib or DMSO-treatment in K562 cells (>900 million contacts) (**Fig. 5A**). PARP1 inhibition by Olaparib decreased the contact frequency between putative enhancer-promoter pairs (*p* < 1×10^−100^; Wilcoxon signed-rank test) (**Fig. 5B-C**). The effect of Olaparib did not affect contacts between CTCF binding sites, indicating that it was specific to enhancer-promoter contacts. (**Figs. 5C and S7A**). Hence, we conclude that PARP1 enzymatic activity has a role in stabilizing enhancer-promoter contacts.

**Figure 5.**
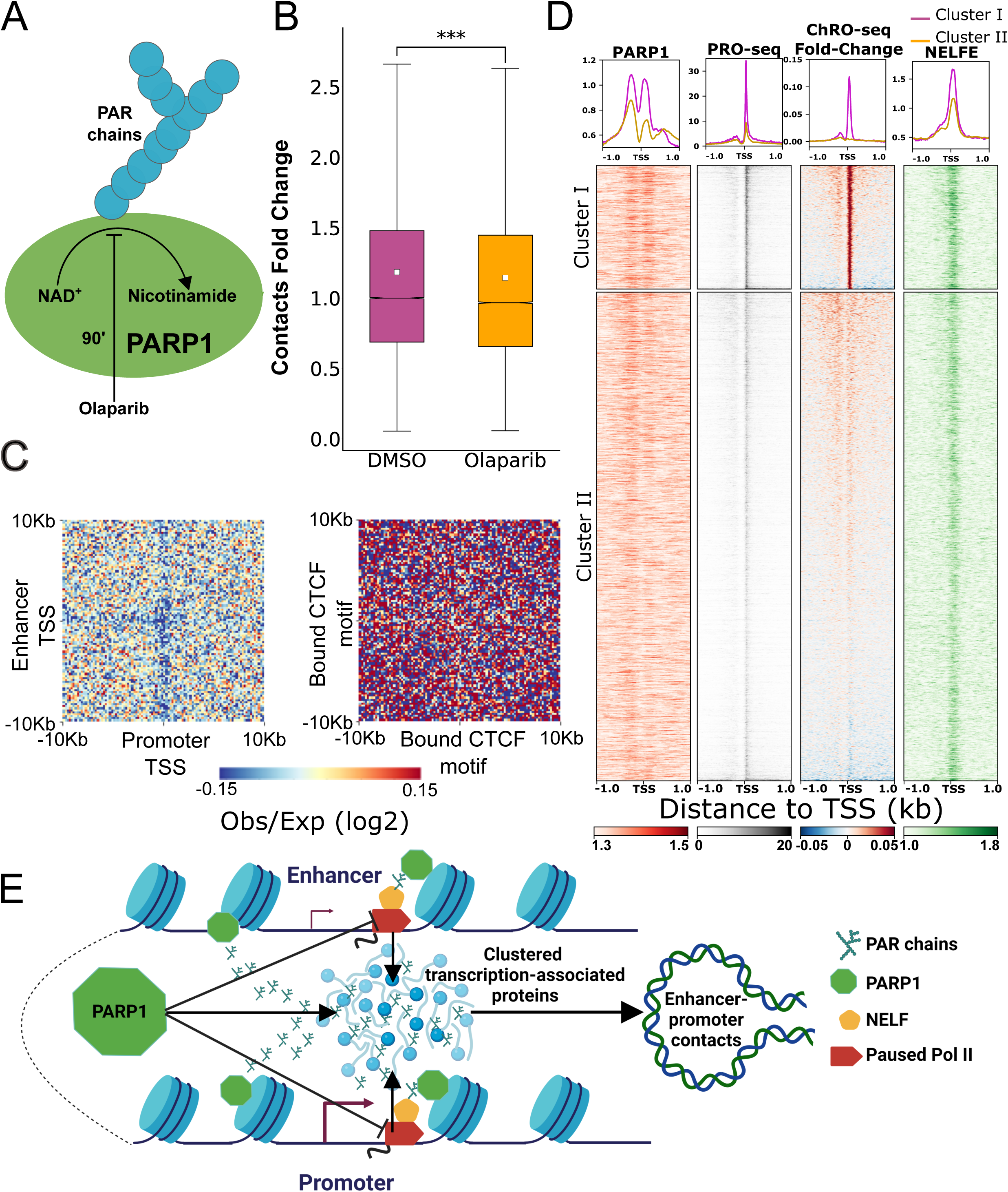
PARP1 inhibition results in a global loss of enhancer-promoter contacts. (A) Illustration of the experiment using Olaparib to inhibit PAR chain formation by PARP1. (B) Boxplot comparing the distribution of enhancer-promoter contacts over background relative to the median of the DMSO control (left). (C) APA heatmap represents the change in contacts compared to DMSO control for K562 cells following PARP1 inhibition for enhancer-promoter contacts (left) and contacts between transcriptionally inactive boud CTCF motifs (right). (D) Heatmaps and metaplots showing signal from PARP1 CUT&Tag, PRO-seq, and NELFE ChIP-seq as well as the fold change in Pol II density following PARP1 inhibition, calculated from ChRO-seq at K562 promoters. Heatmaps are clustered based on the fold-change in ChRO-seq signal following PARP1 inhibition. (E) Model showing the bidirectional effect of PARP1 enzymatic activity on enhancer-promoter contacts: PARP1 catalyzes the formation of PAR chains that, along with paused Pol II, promote the formation of clusters of transcription-associated proteins near enhancer and promoter TSSs. On the other hand, the PARylation of NELFE promotes Pol II escape from pause, reducing local concentrations of paused Pol II and destabilizing enhancer-promoter contacts.

PARP1 promotes the escape of paused Pol II by PARylating NELFE, a subunit of the NELF protein complex (*55*). Indeed, ChRO-seq data in Olaparib treated K562 cells showed a global increase in paused Pol II near TSSs relative to a DMSO control (**Fig. S7B**). To understand the impact of PARP1 on Pol II pausing, we prepared high-resolution PARP1 CUT&Tag (*64*) data in K562 cells. PARP1 CUT&TAG revealed distinct binding patterns of PARP1 in promoter and enhancer TSSs: whereas PARP1 bound at similar levels both upstream and downstream of the TSS at enhancers, PARP1 binding downstream of the TSS varited significantly between different promoters (**Fig. S7B**). At steady state, binding of PARP1 downstream of promoter TSSs was correlated with the abundance of both paused Pol II and NELFE (**Fig. S7B**). Following Olaparib treatment, promoters with a large downstream signal for PARP1 had a substantial increase in paused Pol II (Cluster I; **Fig. 5C**). This additional downstream PARP1 peak, unique to promoters that correlated with increased pausing, is located roughly at the +1 nucleosome, near where paused Pol II and NELFE reside on the chromatin (**Fig. 5C**) (*55*). Notably, the effect of PARP1 inhibition on Pol II signal was strongly positively correlated with baseline levels of PARP1 binding near the TSS (**Fig. S7C**). Collectively, these findings indicate that while PARP1 is located at most active promoter regions, we see substantial variability in the effect of PARP1 on paused Pol II between different promoters.

We reasoned that increased Pol II pausing, which increases enhancer-promoter contact frequency, may partially mask other effects of PARP1 that lead to decreased contact frequency observed following Olaparib. Consistent with this model, we found that changes in paused Pol II density near promoter TSSs were correlated with changes in enhancer-promoter contact frequency (Pearson’s R = 0.30, *p* = 0.003; taking median of 1-percentiles, see Methods), such that candidate enhancer-promoter pairs which accumulated more paused Pol II tended to lose fewer contacts. These findings suggest that PARP1 has at least two effects on enhancer-promoter contacts: The main effect, a stabilizing one, is independent of Pol II pausing and may depend on the contribution of PAR chains to macromolecular clustering (*62*). A weaker, destabilizing effect of PARP1 activity on enhancer-promoter contacts depends on PARylating NELFE (*55*) and accelerating Pol II escape from pause (**Fig. 5E**).

## Discussion

Currently two alternative models are proposed to explain how enhancer and promoter regions communicate. The looping model holds that enhancer and promoter DNA come into close physical contact, perhaps through a structural bridge formed by proteins involved in transcriptional activation (*5, 6*). More recently, an alternative model (which we will call the “hub” model, following (*37, 43*)) has come into favor which predicts that enhancer and promoter regions form malleable “hubs”, in which very high local concentrations of transcription-associated proteins facilitate transcriptional bursts. The hub model was proposed to address several recent findings which did not appear compatible with a loop model, including long physical distance between enhancers and promoters upon gene activation (*16, 17, 65*), the apparent ability of enhancers to activate transcription from multiple promoters simultaneously (*66*), multi-way interactions of enhancer clusters (*67*), and observations that transcription-associated proteins, proposed to compose a structural enhancer-promoter bridge, have modest effects on enhancer-promoter interaction frequency (*20, 22, 46*).

A key difference between the loop and hub models is that the loop model predicts a short physical distance between enhancers and promoters upon interaction. We used the nucleosome-level resolution of Micro-C in conjunction with a large-scale CRISPRi screen to demonstrate that functional pairs of enhancers and promoters are most enriched in contacts near the TSS, with a peak at interactions between the +1 (enhancer)/ +1 (promoter) and the +2/ +2 nucleosomes. Micro-C only detects contacts that are close enough to be crosslinked and ligated (*9*), suggesting that functional enhancer-promoter pairs are enriched for very short interaction distances. Precisely how close enhancer and promoter regions are remains difficult to say (our findings highlight the need for new 3C-based methods that measure distance directly). Although nucleosomes are 10 nm in diameter (*68*) the free ends of DNA sequence are even closer together than 10 nm when they exit the nucleosome at the dyad axis (*69*). This implies that DNA encoding enhancer-promoter elements can come in close proximity at some point during or around transcriptional activation. We caution that our findings do not mean the loop model is correct; rather our findings may alternatively imply that distances between functionally interacting enhancers and promoters may not be as large as previous estimates appear to suggest. We propose that short-distance enhancer-promoter interactions are functional, but are more malleable than predicted by the looping model (*5, 6*).

A key prediction of both models is that transcription-associated proteins, including transcription factors, mediator and Pol II play key roles in enhancer-promoter communication. For this reason, the muted effect that degrading key transcription proteins, including mediator and Pol II, was reported to have on contact frequency was unexpected (*20, 22, 46*). Chromatin at enhancers and promoters undergoes substantial changes when Pol II or other transcription proteins are depleted (*70, 71*), and these changes must be controlled for when measuring changes in contact frequency. Normalizing to the local changes in background contacts highlight Pol II as a major factor that contributes to enhancer-promoter interaction frequency. Notably, our results are consistent with very new preprints that make use of the higher-resolution MNase-based 3C analyses following depletion of Pol II or Mediator (*72, 73*). Several aspects of Pol II may help facilitate interactions, especially under a hub model: First, the C-terminal heptad repeats on RPB1, the largest subunit of Pol II, have been shown to aid in macromolecular clustering (*38, 41, 42, 65, 74, 75*). Second, the nascent RNA emerging from the exit channel may also contribute to clustering (*76, 77*).

We present several independent lines of evidence that highlight a role for paused Pol II in stabilizing enhancer-promoter interactions. Paused Pol II can be stable over durations estimated between 1-10 minutes (*47, 78*). Given its stable attachment to DNA through the transcription bubble, it is possible that paused Pol II may serve as one of the tethers connecting promoter or enhancer DNA into an enhancer-promoter interaction. Under a hub model, paused Pol II initiated from multiple TSSs within a transcription initiation domain (*36*) may serve to keep both enhancer and promoter DNA tethered to the hub. Indeed, paused Pol II tethering enhancers into a hub may serve as one way in which enhancer-templated RNAs (eRNAs) have a sequence-independent biological function. This type of multivalent interaction across multiple Pol II molecules within a narrow window is more difficult to imagine in a loop model, which posits physical interactions between molecules at the enhancer and target promoter.

Our observation that PAR chains catalyzed by PARP1 appear to stabilize enhancer-promoter contacts may be most compatible with a hub model. PAR chains facilitate macromolecular clustering via transient interactions between IDRs (*62, 79*). PARP1 also has an additional role in releasing paused Pol II into productive elongation by parylating NELFE (*55*). Our finding that PARP1 inhibition increased paused Pol II is consistent with this previous report. At first glance, this result appears incompatible with our observations that paused Pol II stabilizes contacts, but that inhibition of PARP1 reduces contacts. The most parsimonious explanation is that PARP1 has multiple effects on enhancer-promoter contacts: some of which stabilize interactions between transcription related proteins, and at least one which reduces contacts by releasing paused Pol II. The correlation observed between increased paused Pol II and contact frequency appears to support this multi-function model.

In summary, our work suggests several important changes to the prevailing models of enhancer-promoter interactions. First, we found that functionally active enhancer-promoter pairs are more likely to reside in close physical proximity. Second, we provide direct evidence for the effect of Pol II on enhancer-promoter contacts. Our work emphasizes an important effect of Pol II pausing in metazoan cells and sheds light on the evolution of this trait alongside long-range enhancer-promoter interactions. Third, we show that PARP1, a major chromatin modulator and transcriptional regulator, affects enhancer-promoter communication in several ways (**Fig. 5**). Thus, considering transcription as a modulator of enhancer-promoter contacts may help future studies to better define the temporal correlation between the two.

## Materials and methods

### Cell culture

Cells were cultured in a humidified 37°C incubator with 5% CO2. K562 and Jurkat cells were grown in RPMI-1640 medium supplemented with 10% fetal bovine serum and 1X penicillin streptomycin antibiotic. For the PARP1 inhibition experiment, cells were treated with either 10 μM Olaparib, initially diluted in DMSO, or DMSO only, for 90 min before immediately crosslinked for Micro-C or lysed in NUN buffer (0.3 M NaCl, 1 M Urea, 1% NP-40, 20 mM HEPES, pH 7.5, 7.5 mM MgCl2, 0.2 mM EDTA, 1 mM DTT, 20 units per ml SUPERase In RNase Inhibitor (Life Technologies, AM2694), 1X Protease Inhibitor Cocktail (Roche, 11 873 580 001)) for ChRO-seq.

mECSs harboring a homozygous endogenous NELFB-FKBP12^F36V^ fusion protein were cultured on 0.1% gelatin (Millipore) in PBS+/+ coated tissue-culture grade plates. For routine culture, cells were grown in Serum/LIF conditions: DMEM (Gibco), supplemented with 2 mM L-glutamine (Gibco), 1x MEM non-essential amino acids (Gibco), 1 mM sodium pyruvate (Gibco), 100 U/ml penicillin and 100 U/ml streptomycin (Gibco), 0.1 mM 2-mercaptoethanol (Gibco), 15% Fetal Bovine Serum (Gibco), and 1000 U/ml of recombinant leukemia inhibitory factor (LIF).

To induce NELFB degradation, dTAG-13 (Bio-Techne) was reconstituted in DMSO (Sigma) at 5 mM. dTAG-13 was diluted in maintenance medium to 500 nM and added to cells with medium changes for the specified amounts of time. For dTAG washes, the cells were washed 4 times, twice with PBS +/+ and twice with maintenance medium following the treatment time to ensure complete removal of the dTAG ligand. At the end of each dTAG-13 treatment time point, cells were detached using Trypsin-EDTA (0.05%) (Gibco) and counted before crosslinking for Micro-C.

### Micro-C

Micro-C for K562, Jurkat and mESCs was performed by following the published protocol for mammalian Micro-C (*21, 22, 80*). Cells were crosslinked with 1 ml per million cells of 1% formaldehyde for 10 minutes at room temperature and quenched by 0.25 M Glycine for 5 min. After spin-down for 5 minutes at 300Xg at 4 °C, cells were washed at a density of 1 ml per million cells in ice cold PBS. Cells were crosslinked a second time, with 1 ml per 4 million cells of 3 mM disuccinimidyl glutarate (DSG) (ThermoFisher Scientific, 20593) for 40 min at room temperature and quenched by 0.4 M Glycine for 5 min. Following two washes with ice cold PBS, cells were flash-frozen and kept at -80°C until further use. For MNase digestion, cells were thawed on ice for 5 min, incubated with 1ml MB#1 buffer (10 mM Tris-HCl, pH 7.5, 50 mM NaCl, 5 mM MgCl2, 1 mM CaCl2, 0.2% NP-40, 1x Roche cOmplete EDTA-free (Roche diagnostics, 04693132001)) and washed twice with MB#1 buffer. MNase concentration for each cell type was predetermined using MNase titration experiments exploring 2.5-20U of MNase per million cells. We selected the MNase concentration that gives ∼90% mononucleosomes. Chromatin was digested with MNase for 10 min at 37 °C and digestion was stopped by adding 8 ul of 500 mM EGTA and incubating at 65 °C for 10 min.

Following dephosphorylation with rSAP (NEB #M0371) and end polishing using T4 PNK (NEB #M0201), DNA polymerase Klenow fragment (NEB #M0210) and biotinylated dATP and dCTP (Jena Bioscience #NU-835-BIO14-S and #NU-809-BIOX-S, respectively), ligation was performed in a final volume of 2.5 ml for 3h at room temperature using T4 DNA ligase (NEB #M0202). Dangling ends were removed by a 5 min incubation with Exonuclease III (NEB #0206) at 37 °C and biotin enrichment was done using 20 ul Dynabeads™ MyOne™ Streptavidin C1 beads (Invitrogen #65001). Libraries were prepared with the NEBNext Ultra II Library Preparation Kit (NEB #E7103). Samples were sequenced on a combination of Illumina’s NovaSeq 6000 and HiSeq 2500 at Novogene.

### Micro-C data mapping and visualization

All Micro-C mapping was done using the mirnylab/distiller-nf: v0.3.3 pipeline (*81*). Raw data were mapped to the hg38 human genome assembly (K562 and Jurkat) or mm10 mouse genome assembly (mESCs). For analysis of contacts in the *MYC* locus, data was mapped to hg19 human genome assembly due to a large gap present in this locus when mapping K562 sequencing data to hg38. For data visualization by contact maps, multi cool (mcool) files, balanced by iterative correction and eigenvector decomposition (ICE) for resolutions of 200 bp to 10 Mb were generated from contacts with both ends having a mapq score > 30. Micro-C data visualization as contact maps in genome-browser shots with available PRO-seq, dREG, CRISPRi and histone marks tracks was done using the HiCExplorer tool (*82*) and pyGenomeTracks (*83*). Virtual 4C tracks were prepared as described previously (*49*). 1D signal near enhancer and promoter TSSs (**Fig. S2**) was calculated based on the distiller-nf output pairs files, filtered for intra-chromosomal with mapq > 30. Contacts assigned to the 5’ of single reads were shifted 75bp downstream, based on their orientation, to the probable center of the nucleosome.

### ChRO-seq

For chromatin isolation from cultured K562 cells, we added 1 ml of 1X NUN buffer and vigorously vortexed the samples for 1 min. An additional 500 µl of NUN buffer was added and samples were vigorously vortexed for an additional 30 s. The samples were then incubated on ice for 30 min with a brief vortex every 10 min and centrifuged at 12,500g for 30 min at 4 °C, after which the NUN buffer was removed from the chromatin pellet. The chromatin pellet was washed 3 times with 1 ml of 50 mM Tris-HCl, pH 7.5, supplemented with 40 U/ml of RNase inhibitor, centrifuged at 10,000Xg for 5 min at 4 °C, and the supernatant discarded. After washing, 100 µl of chromatin storage buffer (50 mM Tris-HCl, pH 8.0, 25% glycerol, 5 mM Mg(CH3COO)2, 0.1 mM EDTA, 5 mM DTT, 40 U/ml Rnase inhibitor) was added to each sample. The samples were loaded into a Bioruptor and sonicated with the power setting on high, with a cycle time of 10 min with cycle durations of 30 s on and 30 s off. The sonication was repeated up to three times, as needed, to get the chromatin pellet into suspension. Samples were stored at − 80 °C.

ChRO-seq library preparation was performed following a published protocol (*24*), with minor modifications. Chromatin from 1 million K562 cells in 25 µl chromatin storage buffer was mixed with 25 µl of 2X chromatin run-on buffer (10 mM Tris-HCl, pH 8.0, 5 mM MgCl2,1 mM DTT, 300 mM KCl, 10 μM Biotin-11-ATP (Perkin Elmer, NEL544001EA), 40 μM Biotin-11-CTP (Perkin Elmer, NEL542001EA), 10 μM Biotin-11-GTP (Perkin Elmer, NEL545001EA), 40 μM Biotin-11-UTP (Perkin Elmer, NEL543001EA), 2ng/μl Yeast tRNA (VWR, 80054-306), 0.8 U/μl RNase inhibitor, 1% Sarkosyl (Fisher Scientific, AC612075000)). The run-on reaction was incubated in a thermomixer at 37 °C for 5 min at 750 RPM. The reaction was stopped by adding 300 μl of Trizol LS (Life Technologies, 10296-010). To clean up the reaction prior to base hydrolysis, 40 μl of 1-bromo-3-chloropropane (BCP) (Sigma, B9673) was added to the samples and samples were vortexed for 20 sec, incubated for 3 min at RT and centrifuged at 17,000Xg at 4°C for 5 min. ∼250 μl of aqueous phase was transferred to a new 1.5 ml tube and samples were pelleted at 17,000Xg at 4°C for 15 min in 650 ml ice-cold 100% EtOH with GlycoBlue (2.5 μl) (Ambion, AM9515) to visualize the RNA pellet. The pellet was further washed with 0.5 ml ice-cold 75% by vortex-mixing and centrifugation at 17,000Xg at 4°C for 5 min. Any residual EtOH was removed by air-drying the pellet for 5 min in RT. The RNA pellet was resuspended in 20 μl of diethylpyrocarbonate (DEPC)-treated water and heat denatured at 65 °C for 40 sec. For base hydrolysis, 5 μl of 1N NaOH was added to the RNA sample to get 0.2N NaOH, and samples were incubated on ice for 8 min. Base hydrolysis was stopped by adding 25 μl of Tris-HCl pH 6.8 and gently mixing. 3′ Adapter ligation was done using T4 RNA Ligase 1 (NEB, M0204L). A first binding to streptavidin beads (NEB, S1421S) and washed as described (*24*). RNA was removed from beads by Trizol (Life Technologies, 15596-026) and followed by a 5′ decapping with RNA 5′ pyrophosphohydrolase (RppH, NEB, M0356S). The 5′ end was phosphorylated using T4 polynucleotide kinase (NEB, M0201L). A second bead binding was performed, followed by an on-beads 5′ adapter ligation. After bead washes, RNA was removed from beads by adding 300 μl of TRI reagent (MRC, TR 118). 40 μl of BCP were added to the samples and samples were vortexed for 20 sec, incubated for 3 min at room temperature and centrifuged at 17,000Xg at 4°C for 5 min. ∼180μl of aqueous phase was transferred to a new 1.5 ml tube and samples were pelleted at 17,000Xg at 4 °C for 15 min in 450 ml ice-cold 100% EtOH with GlycoBlue (2.5 μl) to visualize the RNA pellet. The pellet was further washed with 0.4 ml ice-cold 75% EtOH by vortex-mixing and centrifugation at 17,000Xg at 4 °C for 5 min. Any residual EtOH was removed by air-drying the pellet for 5 min in RT. A reverse transcription reaction using Superscript III Reverse Transcriptase (Life Technologies, 18080-044) was used to generate cDNA. cDNA was then amplified using Q5 High-Fidelity DNA Polymerase (NEB, M0491L) to generate the ChRO-seq libraries following the protocol provided by NEB. Libraries were sequenced using Illumina’s HiSeq 4000 at Novogene.

### ChRO-seq, PRO-seq and GRO-seq data processing and analysis

Processing for ChRO-seq data (as well as PRO-seq and GRO-seq available raw data) in this study was done using the Proseq2.0 pipeline available from GitHub (https://github.com/Danko-Lab/proseq2.0) (*84*). Differential expression analyses between K562 and Jurkat cells for pausing signal and gene body transcription levels was performed by DEseq2 (*85*) either on signal between the TSS and 250bp downstream (pause signal) or signal downstream to the first 250 bp through the annotated (GENCODE V29) polyadenylation cleavage site (gene body signal). For visualization of the changes, fold change in expression following NELFB-dTAG in mESCs or PARP1 inhibition in K562 was performed by deepTools bigwigCompare command at 1bp resolution, using 0.25 as pseudocount. For the NELFB-dTAG PRO-seq visualization, fold-change and normalized PRO-seq signal matrices were calculated in a stranded manner, followed by a contacennation of the two strands’ matrices to generate a single, stranded matrix (**Fig. 4B**).

### Definition of TIRs, enhancers, promoters and transcription start sites

For mESCs we first defined TIRs genome-wide as detected by dREG (*30, 31*) using available GRO-seq data from mESCs (*47, 86*). To finely and unbiasedly define the position of transcription initiation at each of these TIRs, we used the position with the most 5’ mapped GRO-seq reads within the dREG peak (maxTSN). For the analyses of K562 cells, we first called TIRs using dREG from available PRO-seq data (*31*). The center of these TIRs was defined as the center of enhancers and promoters for the analysis comparison of contacts between functional and nonfunctional enhancer-promoter pairs, based on CRISPRi data. For any further analyses the center of enhancers and promoters was defined as the maxTSNs, called using the data from coPRO with enrichment for 5’ capping (coPRO-capped) (*36*). For the comparison between K562 and Jurkat cell lines, we called TIRs in both K562 and Jurkat using PRO-seq data (*31*) and dREG and determined maxTSN based on coPRO-capped from K562 cells. We defined promoters based on the existence of any known human (K562 and Jurkat) of mouse (mESCs) stable 5’ mapped transcripts from CAGE (*87*) within 5kb away in the direction of maximum initiation. In analyses including Jurkat and K562 cell lines, we considered only shared promoters based on proximity to the best transcription start site defined by the deconvolution of expression for nascent RNA-sequencing data (DENR), based on GENCODE V29 annotations, in both cell types. We used a combined set of enhancers from TIRs detected in both cell lines to define enhancers. Since promoters make a relatively small fraction of all TIRs found in the data and can act as enhancers for other distal genes(*88*) we included promoters under the definition for enhancers whenever we calculated enhancer-promoter contacts genome-wide.

### Micro-C contact normalization and comparison between CRISPRi-defined functional and nonfunctional pairs

Comparison between functional and nonfunctional enhancer-promoter pairs was based either on CRISPRi genetic screens for enhancer function either in the *MYC* locus, based on cell viability (*28*) or based on expression from single-cell RNA sequencing analysis (*27*). In addition to the seven active enhancers in the K562 *MYC* locus, we also detected 54 other TIRs that were marked by DNase-I hypersensitivity sites (DHSs) and histone modifications, were located in the same TAD, and were tested by CRISPRi, but which did not affect the growth rate of K562 cells. These TIRs were considered nonfunctional and compared to the seven functional *MYC* enhancers. For the genome wide-analysis based on data from (*27*), we defined functional pairs as pairs where CRISPR inhibition of the enhancer resulted in a loss of a minimum of 10% of gene expression, with an empirical p-value < 0.05. nonfunctional pairs were defined as pairs with less of 5% effect on gene expression and empirical p-value > 0.9. To remove possible confounding effect, we filtered the functional and nonfunctional pairs to have similar distributions of enhancer-promoter contacts (limited between 5kb and 1Mb), accessibility (by ATAC-seq) and gene body transcription levels (by PRO-seq) in the target gene (**Fig. S1C**). All CRISPRi-targeted enhancers and target promoters were reassigned to their nearest dREG peak center, within 5kb, on the same strand.

Enhancer-promoter contacts were defined as contacts that map to a 5kb window near the promoter on one end and the enhancer on the other hand. Contacts were normalized to the expected based on a non-parametric LOWESS smoothing of the contacts-by-distance function in a region corresponding to a 1Mb in the orientation of the promoter, relative to the enhancer (**Fig. S1A**). Observed over expected ratios were then compared between functional and nonfunctional pairs (**Fig. 1B,C, Fig. S1B**). Differences in contacts between CRISPRi-defined functional and nonfunctional pairs were calculated based on cell-by-cell differences between APA matrices for all functional and all nonfunctional pairs, normalized for the number of pairs. The differences were calculated as the medians of the differences based on 1000 bootstrapping iterations of the functional and nonfunctional pairs, to remove outlier background. These differences were presented as the number of contact differences per 1000 pairs (**Fig. 1E and F**). The APA matrices were centered on the coPRO-based maxTSN as the TSS assigned for each TIR.

### Aggregated Peak Analysis (APA) and APA matrices normalization for comparison between samples

We expected significant changes in chromatin after manipulating Pol II transcription (*70, 71*). As such, not only are enhancer-promoter contacts expected to change, but the background contacts with at least one end originating at enhancer- and promoter-regions may be affected between conditions. As APAs are often used to characterize contacts (*7, 22, 46*), we devised an APA that normalizes enhancer-promoter contacts to all contacts associated with enhancers and promoters. We calculated changes in aggregated 1D signal mapped to the same windows around enhancers and promoters, as used in the APA. We calculated the expected change in each pixel based on the ratios between the sample-specific sum of the 1D signal in each treatment condition (**Fig. 2A**). For all other, intra-sample APA analysis we used the aggregated raw counts. For APAs calculated at windows of 20kb around the anchors, we considered all possible anchor pairs within a genomic distance of 25-150kb. For higher resolution APAs with 2kb window around enhancer and promoter TSSs (**Fig. 1D and Fig. S3**), we considered all possible enhancer-promoter pairs within a genomic distance of 5-100kb.

### Individual contact comparison between samples and treatments

We also devised an alternative normalization scheme which compares the number of contacts between enhancer-promoter pairs to the local background near each enhancer and promoter anchor. We calculated the number of contacts between each pair of anchors (enhancer-promoter or CTCF binding sites) using a 5kb window around each anchor. As a background, we counted the number of contacts between each anchor (in a 5kb window) and regions 10-150 kb from the second anchor (**Fig. 2C**). The ratio between the anchor-to-anchor contacts and background contacts was presented in scatterplots or used to calculate the statistical significance of changes between treatments and samples. To avoid the impact of noise, we analyzed only contacts that met a minimum baseline of anchor-to-anchor contacts (at least 8 contacts per billion contacts (CPB)) in one of the treatment conditions. Since TSS calling data (PRO-seq and coPRO-capped) was more abundant for K562 than Jurkat, when comparing K562 and Jurkat libraries we considered enhancer-promoter pairs with at least 8 CPB in both cell lines, to avoid ascertainment bias. The distribution of ratios between enhancer-promoter and background contacts in treated samples (Olaparib\TRP\FLV or dTAG treated cells) was compared to the median ratio in the respective control samples (**Figs 2D-E, 4C-D, 5B and S6A**). For comparison between cell lines, enhancer-promoter contacts at promoters with increased gene body transcription and\or Pol II pausing signal in one cell line, were compared to their median at the other cell line (**Figs. 3D-E and S4A,C**).

### Definition of CTCF binding sites

Contacts between CTCF binding sites were used as a control to determine whether the effects of a treatment were specific to enhancer-promoter contacts. We defined pairs of CTCF binding sites as CTCF motifs that were shown to bind CTCF based on ENCODE ChIP-seq data, within the same minimum and maximum allowed genomic distances as for enhancers and promoters. We focused only on CTCF sites that show no overlap with any dREG-defined TIR within 5kb.

### PARP1 CUT&Tag

24h prior to the experiment, cells were split at a final concentration of 0.5 million cells/mL in fresh RPMI supplemented with 10% FBS and 1X penicillin streptomycin antibiotic. PARP1 CUT&Tag was done in two replicates with 250,000 K562 cells each for baseline control and anti-PARP1 CUT&Tag. Experiments were done according to the bench top CUT&Tag V.3 protocol (*89*), using CUTANA™ Concanavalin A Conjugated Paramagnetic Beads (EpiCypher, SKU: 21-1401) and pAG-Tn5 for CUT&Tag (EpiCypher, SKU: 15-1017), with the following exceptions: (A) cells were not cross-linked and (B) NE1 buffer was used to extract and permeabilize nuclei. For anti-PARP1 CUT&Tag, we used an antibody against the N-terminal of PARP1 (Active Motif, AB_2793257). An anti-IgG antibody (Cell signaling, Normal Rabbit IgG #2729) was used as baseline control.

### CUT&Tag data processing

fastq files were trimmed from Nextera adaptor-associated sequences using cutadapt (*90*) and mapped to hg38 human genome assembly using bowtie2 (*91*). We then used bedtools genomecov to generate bedgraph files and calculated background-normalized PARP1 binding using macs2 bdgcmp command (*92*).

### Statistical analysis

Throughout the manuscript, Mann-Whitney U test is used for independent samples, such as comparison of changes between different sets of genomic loci or pairs. Wilcoxon signed-rank test is used for paired samples, usually being the same loci\pairs compared between samples\conditions. For assessment of trends in our data, such as the changes in the difference in contacts between functional and nonfunctional pairs, or assessing the effect of changes in paused Pol II occupancy on changes in enhancer-promoter contacts, we used Pearson’s correlation coefficient (R). The confidence intervals for the medians throughout the manuscript were calculated using 1000 iterations of bootstrap.

## Data and code availability

Micro-C, CUT&Tag and ChRO-seq data generated in this study were deposited in the Gene Expression Omnibus (GEO) database under accession number GSE206133. H3K27ac and H3K4me2 ChIP-seq data from K562 cells were downloaded from GSE163043. K562 data for ATAC-seq (ENCSR868FGK), CTCF ChIP-seq (ENCSR447BSF), MNase-seq (ENCSR000CXQ) and NELFE ChIP-seq (ENCSR000DOF) were downloaded from ENCODE. DMSO, TRP and FLV treated mESCs Micro-C data (*22*) were downloaded from GSE130275. PRO-seq data (*30, 33*) for Jurkat T-cells were downloaded from GSE66031 and for K562 from GSE60455. PRO-seq data for mECSs harboring a homozygous endogenous NELFB-FKBP12^F36V^ fusion protein, treated and untreated with dTAG-13 (*93*), were downloaded from GSE196653. GRO-seq data for mESCs (*47, 86*) were downloaded from GSE43390 and GSE48895. Positions for human (hg38) and mouse (mm10) CAGE peaks were downloaded from the FANTOM5 database (https://fantom.gsc.riken.jp/5/). All data normalization and visualization code is available at https://github.com/Danko-Lab/E-P_contacts.

## Acknowledgements

We thank E. Apostolou and members of her lab for commenting on a manuscript draft as well as members of the Danko, Lis, and Yu labs for valuable discussions and suggestions throughout the life of this project. Work in this publication was supported by R01-HG010346 and R01-HG009309 (NHGRI) to CGD. The content is solely the responsibility of the authors and does not necessarily represent the official views of the US National Institutes of Health. Some of the figures in this manuscript were created using BioRender.

## Figure legends

**Figure S1.**
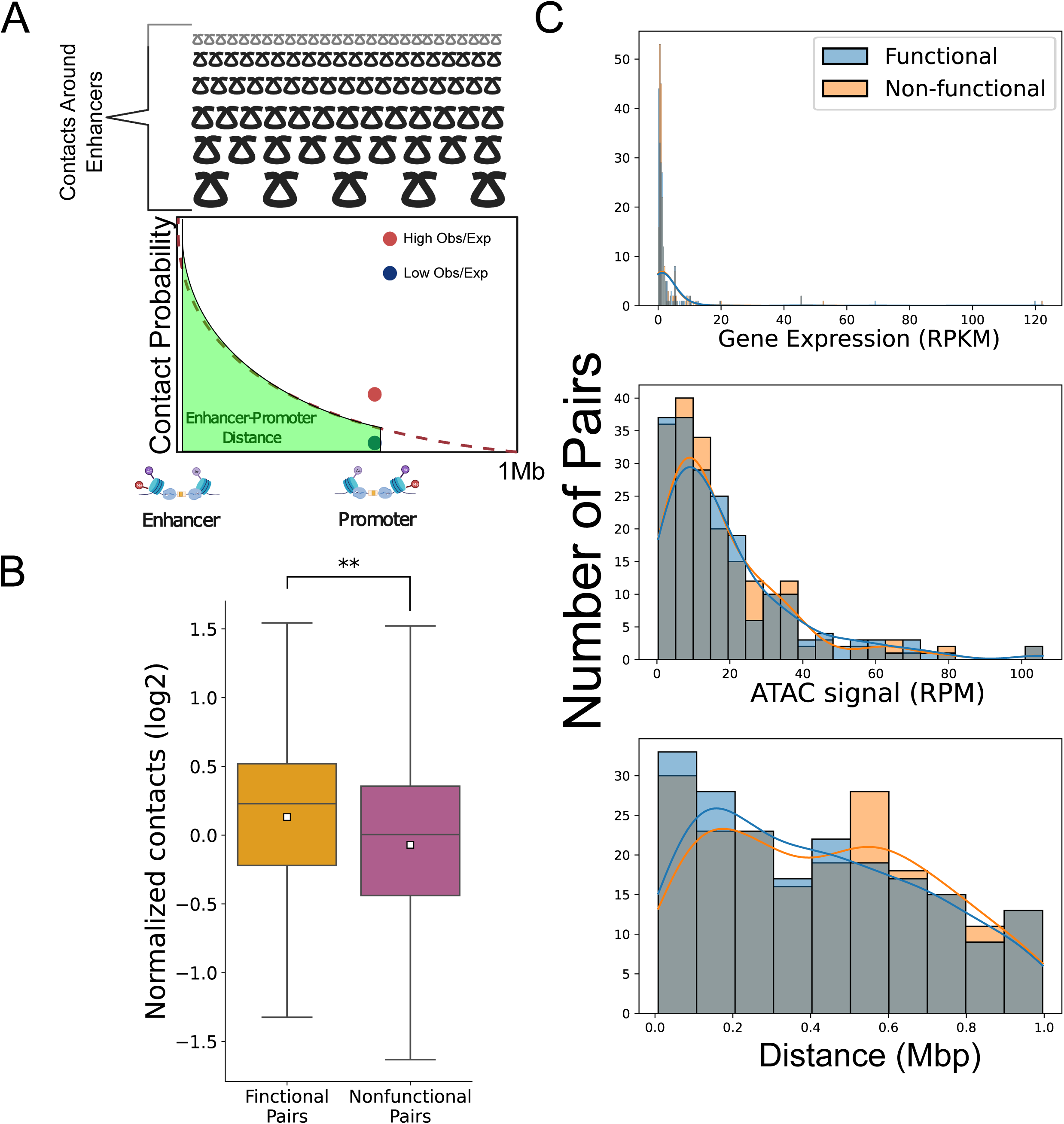
Supplementary details for the comparison between functional and nonfunctional enhancer-promoter pairs. (A) Schematic representation of the LOWESS-based normalization for enhancer-promoter contacts. (B) Box-plot comparing contact levels relative to expected by local distance-decay function of functional versus the nonfunctional enhancer-promoter pairs in the genome, before matching for enhancer-promoter distance, accessibility or target gene expression (** Mann-Whitney p-value < 1×10^−10^). (C) Histograms demonstrating the distribution of functional and nonfunctional enhancer-promoter pairs in terms of PRO-seq target gene transcription signal in reads per kilobase per million reads (RPKM) (top), accessibility by mean ATAC-seq signal (middle) and enhancer-promoter genomic distance (bottom), after matching for these confounding effects.

**Figure S2.**
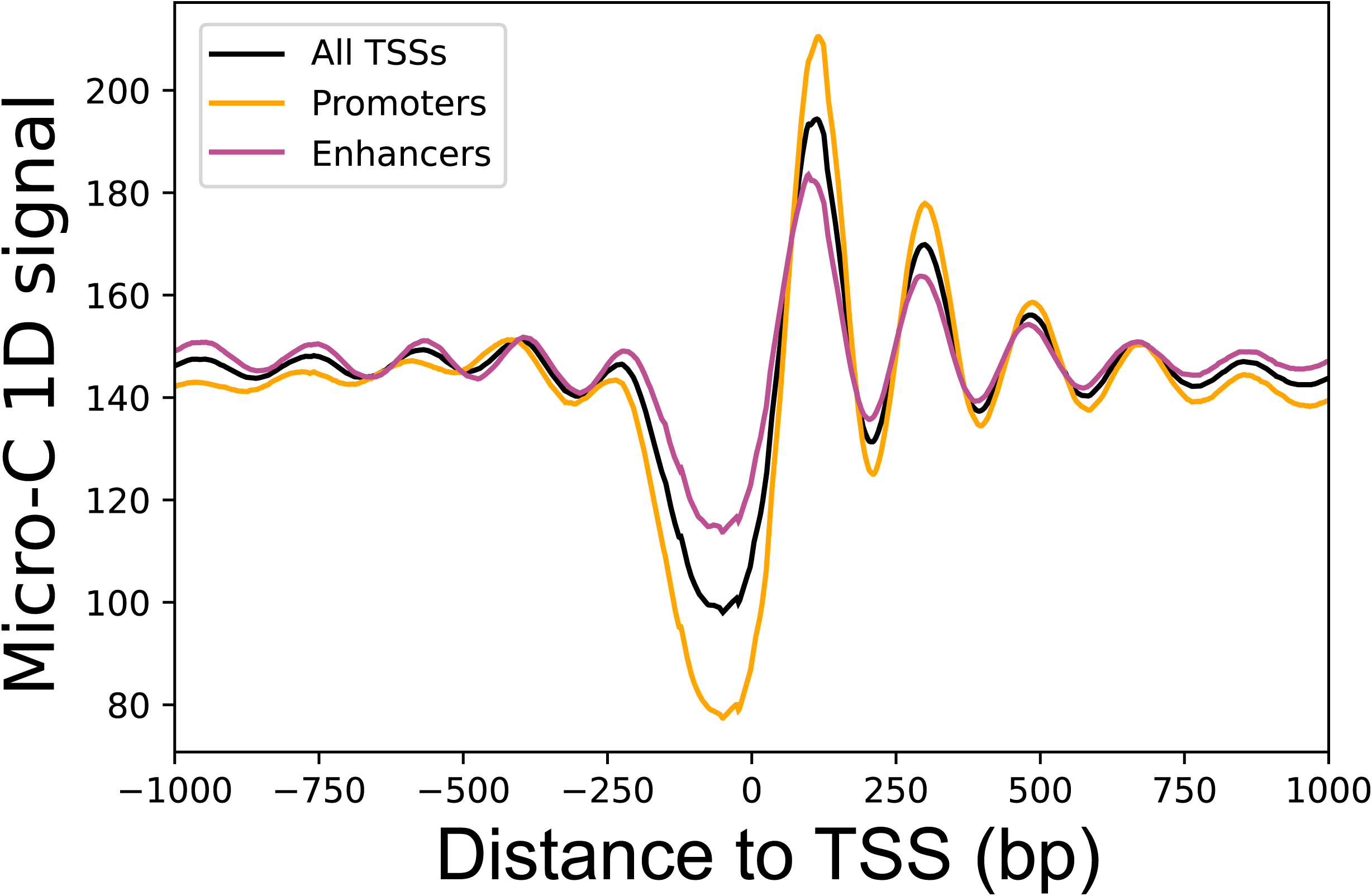
Micro-C 1D signal near TSS genome-wide. One dimensional contact signal for intra-chromosomal contacts with both sides having mapping quality (mapq) ≥ 30. Total median signal was smoothed using a sliding window of 100bp. Shown are signals around promoter TSSs (orange), enhancer TSSs (purple) and all TSSs genome-wide (black).

**Figure S3.**
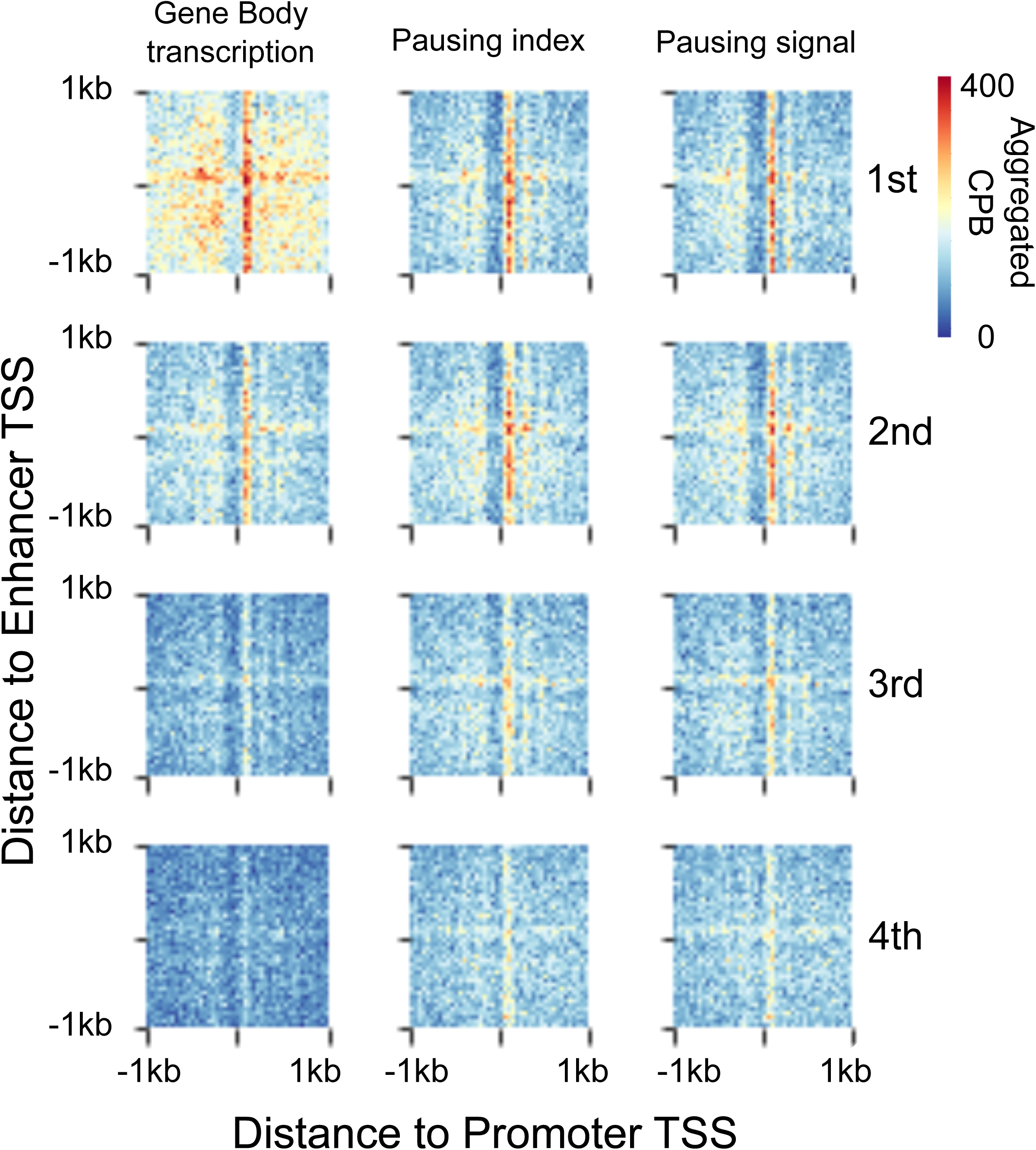
High resolution representation of changes in enhancer-promoter contact architecture associated with gene body transcription and pausing at promoters. APA maps showing enhancer-promoter contacts associated with promoters of the four gene body transcription (left), pausing index (middle) and pause peak signal (right) quartiles. Color Scale represents contacts per billion contacts (CPB). Pixel size is 20bp square.

**Figure S4.**
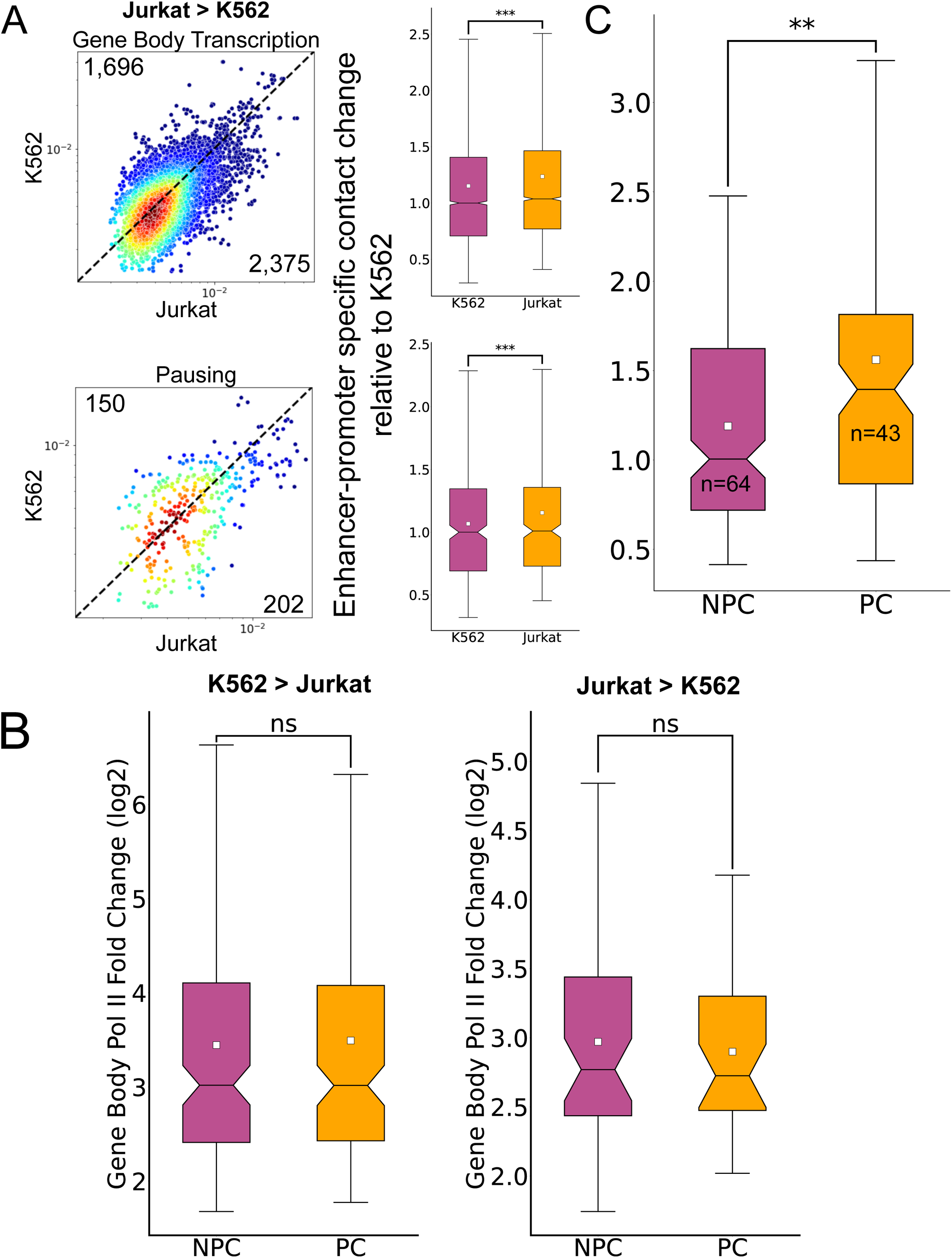
Distribution of fold change in gene body transcription for K562 and Jurkat upregulated genes. (A) Scatterplots and boxplots demonstrating changes in enhancer-promoter contacts associated with promoters of genes with significantly higher gene body transcription (top) and pausing signal (bottom) at Jurkat T-cells compared to K562 (*** Wilcoxon signed-rank test p-value < 1×10^−100^). (B) Boxplots showing the distributions of fold change in gene body signal in genes with no associated paused Pol II change (NPC) and associated significant paused Pol II change (PC) (“ns” - Mann-Whitney p-value > 0.5). (C) Boxplot depicting the relative increase of enhancer-promoter contacts associated with promoters of genes with upregulated gene body transcription in Jurkat T-cells with a corresponding significant increase in pausing signal (pause change - PC) and without a change in pausing signal (no pause change - NPC). (** Mann-Whitney p-value < 1×10^−10^)

**Figure S5.**
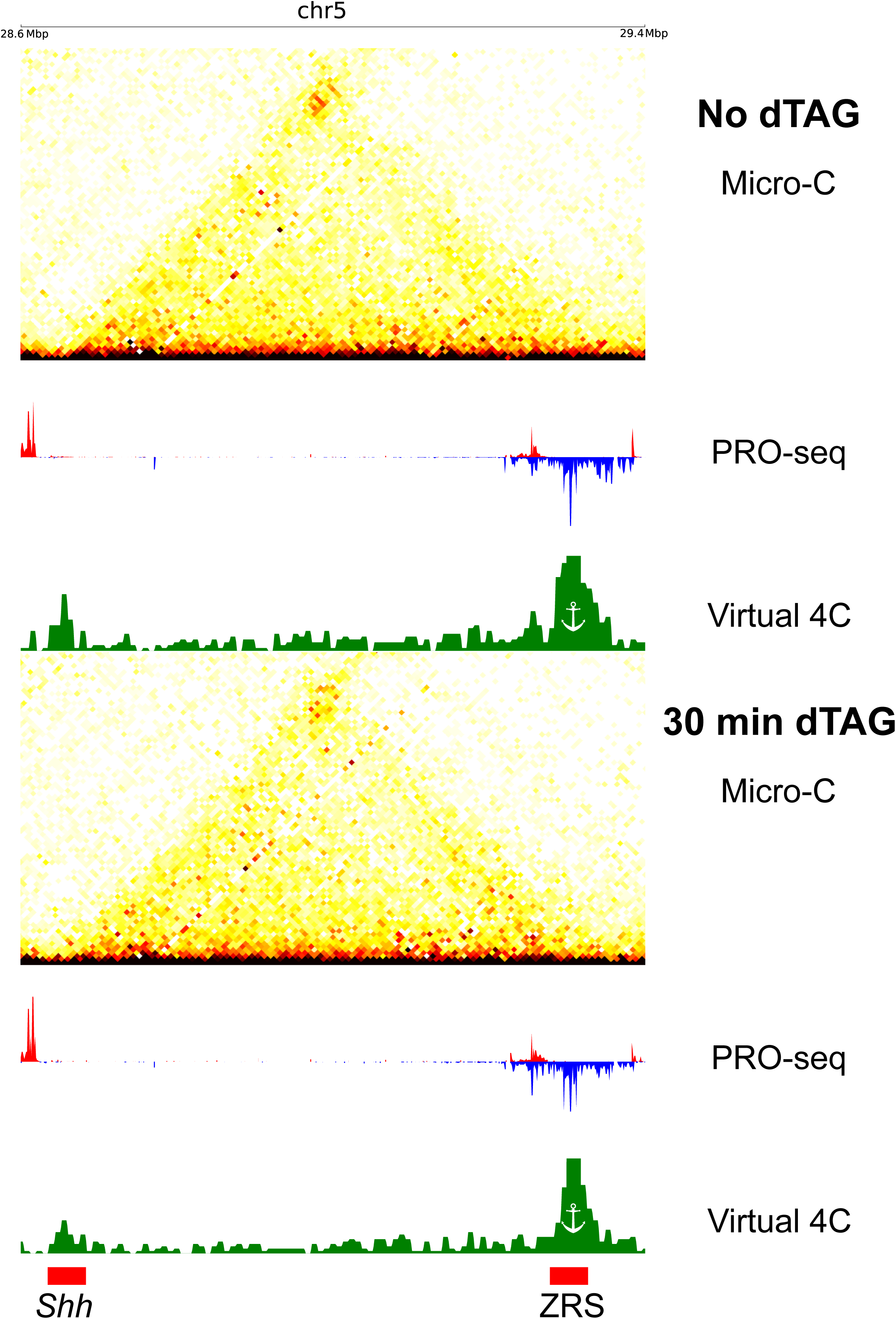
Changes in ZRS-*Shh* contacts following NELFB depletion. Micro-C contact maps in 10kb resolution along with the associated virtual 4C signal and PRO-seq signal in mESCs not treated (top) or treated (bottom) with the dTAG ligand for 30 minutes to degrade NELFB. The positions of the ZRS enhancer and the *Shh* promoter are indicated in red rectangles.

**Figure S6.**
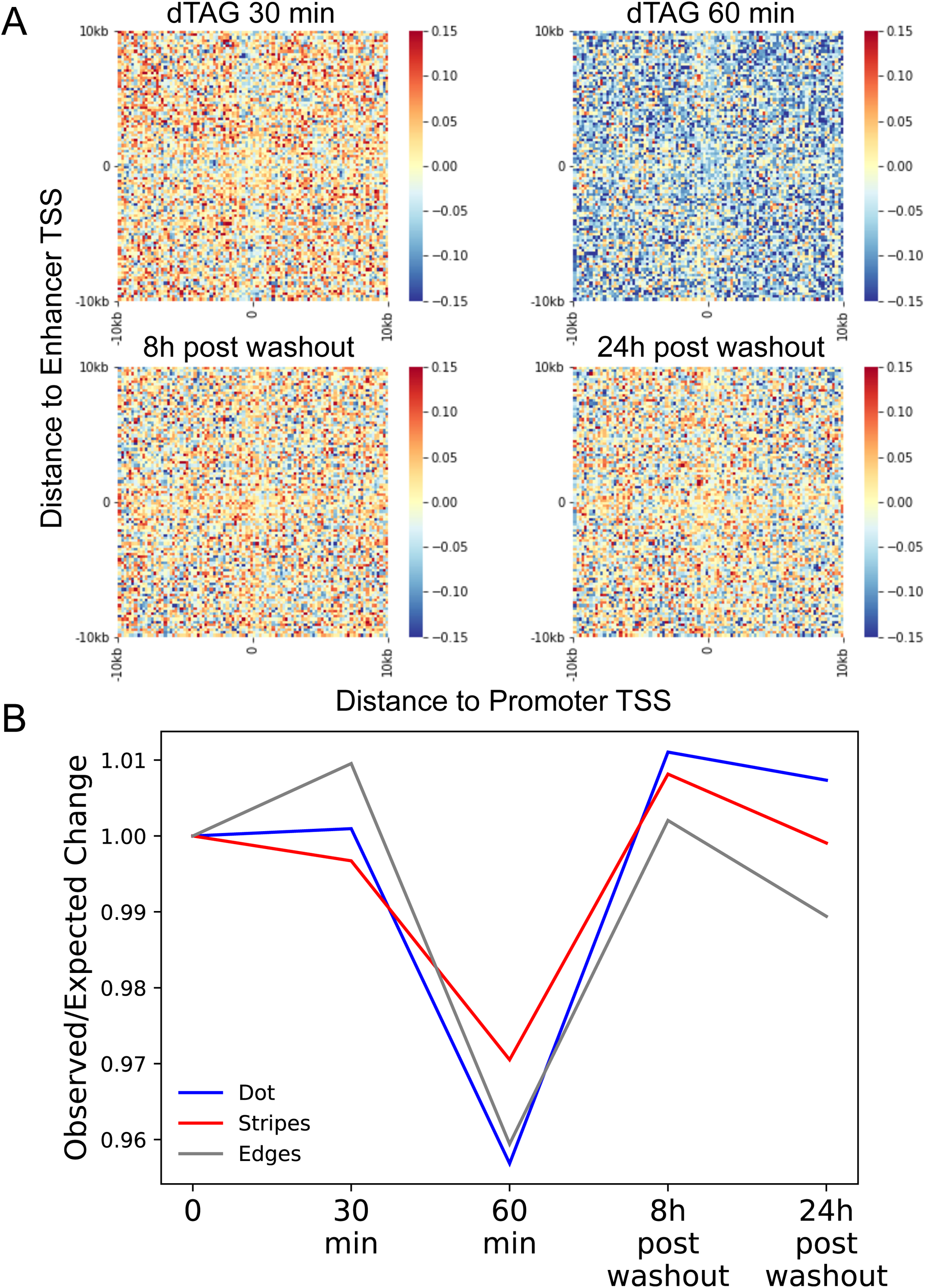
Changes in enhance-promoter contacts architecture following NELFB depletion. (A) APA maps of the observed over expected (log2) changes in enhancer-promoter contacts at 20kb around TSSs. Pixel size is 200bp square. (B) Line plot of the median observed/expected changes at the dot (blue), stripes (red) and edges (gray) relative to T=0 at the different time points of dTAG treatments and following dTAG washout.

**Figure S7.**
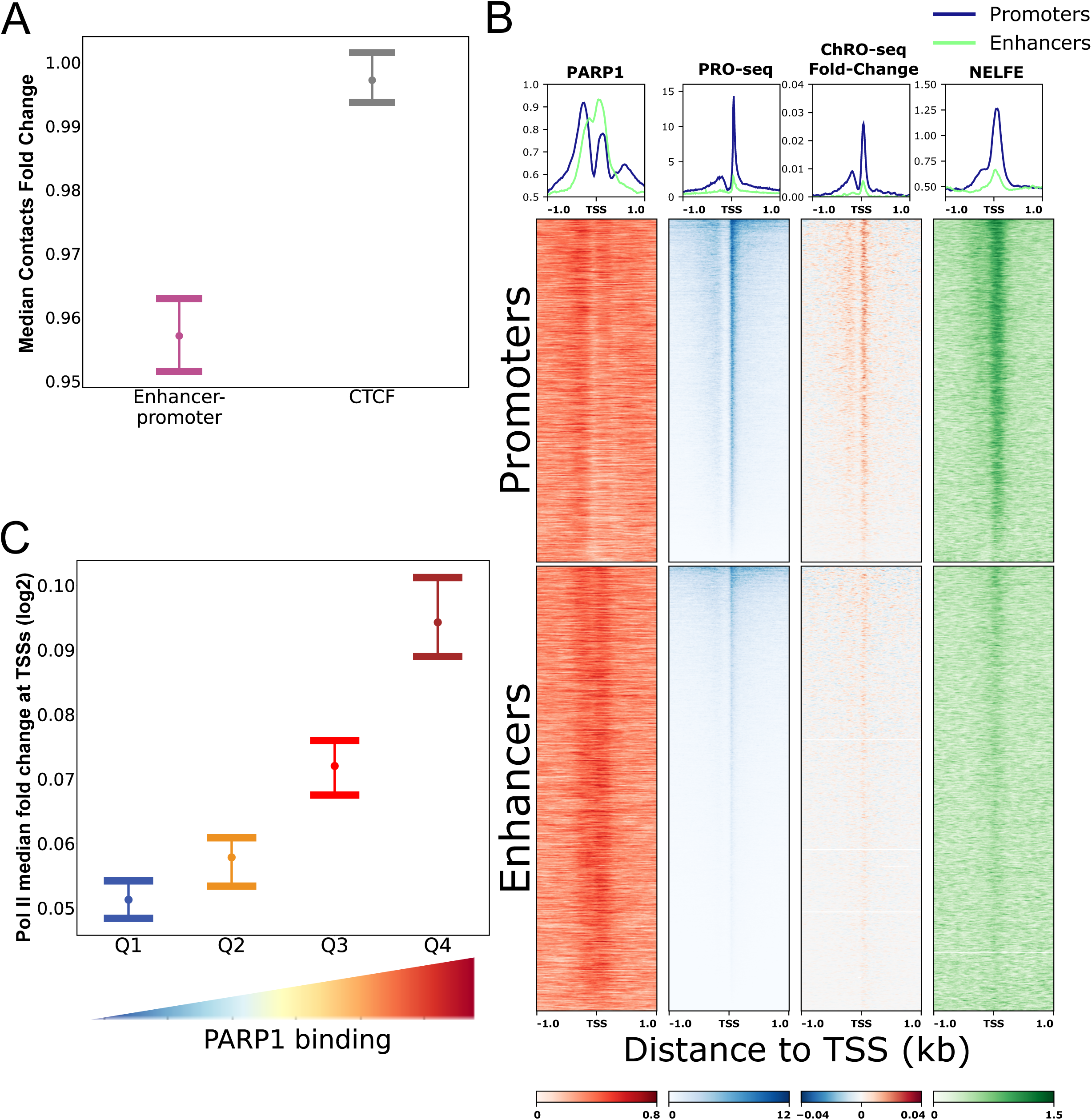
The effect of PARP1 inhibition on enhancer-promoter contacts and transcription at enhancers and promoters. (A) Dot and error plot demonstrating the median contact change and the 95% confidence interval of the median based on 1000 bootstrap iterations for enhancer-promoter (purple) and transcriptionally inactive CTCF motif contacts (gray), following PARP1 inhibition. (B) Heatmaps and metaplots showing signal from PARP1 CUT&Tag, PRO-seq, and NELFE ChIP-seq as well as the fold change in Pol II density following PARP1 inhibition, calculated from ChRO-seq at promoter (top) and enhancer (bottom) TSSs. (C) Dot and error plot demonstrating the median contact change and the 95% confidence interval of the median based on 1000 bootstrap iterations for mean Pol II fold change (log 2 scale) in the four quartiles of mean PARP1 binding in pairs of enhancers and promoters.

